# Cluster similarity spectrum integration of single-cell genomics data

**DOI:** 10.1101/2020.02.27.968560

**Authors:** Zhisong He, Agnieska Brazovskaja, Sebastian Ebert, J. Gray Camp, Barbara Treutlein

## Abstract

Technologies to sequence the transcriptome, genome or epigenome from thousands of single cells in an experiment provide extraordinary resolution into the molecular states present within a complex biological system at any given moment. However, it is a major challenge to integrate single-cell sequencing data across experiments, conditions, batches, timepoints and other technical considerations. New computational methods are required that can integrate samples while simultaneously preserving biological information. Here, we propose an unsupervised reference-free data representation, Cluster Similarity Spectrum (CSS), where each cell is represented by its similarities to clusters independently identified across samples. We show that CSS can be used to assess cellular heterogeneity and enable reconstruction of differentiation trajectories from cerebral organoid single-cell transcriptomic data, and to integrate data across experimental conditions and human individuals. We compare CSS to other integration algorithms and show that it can outperform other methods in certain integration scenarios. We also show that CSS allows projection of single-cell genomic data of different modalities to the CSS-represented reference atlas for visualization and cell type identity prediction. In summary, CSS provides a straightforward and powerful approach to understand and integrate challenging single-cell multi-omic data.

## Background

Recent advances in molecular, engineering, and sequencing technologies have enabled the high-throughput measurement of transcriptomes and other genomic features in thousands of single cells in a single experiment [1–4]. Single-cell RNA sequencing (scRNA-seq) greatly enhances our capacity to resolve the heterogeneity of cell types and cell states in biological samples, as well as to understand how systems change during dynamic processes such as development. However, current scRNA-seq technologies only provide molecular snapshots of a limited number of measured samples at a time. Joint analysis on many samples across multiple experiments and conditions is often required. In such a scenario, the biological variation of interest is usually confounded by other factors, including sample sources and experimental batches. This is particularly challenging for developing systems, where cell states coexist at different points along various differentiation trajectories such as mature cell types as well as intermediate states. Several computational integration methods, including but not limited to MNN [5], Seurat [6, 7], Harmony [8], LIGER [9], Scanorama [10], and Reference Similarity Spectrum (RSS) [11, 12] have been developed to address some of these issues. Among them, MNN identifies mutual nearest neighbors between two data sets and derives cell-specific batch-correction vectors for integration. Seurat corrects for batch effects by introducing an anchoring strategy, with anchors between samples defined by canonical correlation analysis. Harmony uses an iterative clustering-correction procedure based on soft clustering to correct for sample differences. LIGER adapts integrative non-negative matrix factorization to identify shared and dataset-specific factors for joint analysis. Scanorama generalizes mutual nearest-neighbors matching in two data sets to identify similar elements in multiple data sets in order to support integration of more than two data sets. RSS achieves integration by representing each cell by its transcriptome’s similarity to a series of reference samples. Benchmarking on these integration methods have revealed varying performance of each method based on the given scenario highlighting that there is no single magic bullet capable of always dissecting out meaningful variation of interest [13].

Here, we propose an unsupervised scRNA-seq data representation namely Cluster Similarity Spectrum or CSS, which enables integration of single-cell genomic data. Instead of using external references as in RSS, CSS considers every cell cluster in each sample as an intrinsic reference for integration, and represents each cell by its transcriptome’s similarity to clusters across samples. The underlying hypothesis of both RSS and CSS is that the undesired confounding factors, e.g. read coverage of different cells, introduce random perturbations to the observed transcriptomic measures which are not correlated with cell type or cell state identities. Once similarities to different references are normalized, the global differences among cells introduced by the random perturbation are neutralized. Afterwards, cells of the same identity principally share similar patterns of normalized similarities. We refer to this pattern as the similarity spectrum. Additionally, as CSS considers all clusters in different samples as references, normalization is done separately across clusters of different samples, so that global differences across samples can be largely eliminated. We apply CSS to various scenarios focusing on data generated from cerebral organoids derived from human induced pluripotent stem cells (iPSCs). In addition, we also apply CSS to scRNA-seq data sets of other systems, including peripheral blood mononuclear cells (PBMCs) and developing human retina. We use CSS to integrate data from different iPSC lines, human individuals, batches, modalities, and conditions. We show that technical variation caused by experimental conditions or protocols can be largely reduced with the CSS representation, and CSS has similar or even better performance compared to other integration methods including Scanorama, MNN, Harmony, Seurat v3 and LIGER, which were highlighted in previous benchmarking efforts [13, 14]. We show that CSS also allows projection of new data, either scRNA-seq or scATAC-seq, to the CSS-represented scRNA-seq reference atlas for visualization and cell type identity prediction. The CSS codes are available at https://github.com/quadbiolab/simspec.

## Results

### CSS integrates scRNA-seq data from different organoids, batches and human individuals

To calculate the CSS representation, clustering is first performed on the single-cell transcriptomic data of each sample separately, and average expression profiles are calculated for each cluster (Fig. 1, Supplementary Fig. 1). Transcriptome similarity, here represented as the Spearman or Pearson correlation between gene expression profiles, is then calculated between each single cell and each cell cluster average. For each cell, the calculated similarities are normalized across clusters of each sample and concatenated, resulting in its CSS representation. Different normalization methods can be used, including z-transformation and kernel probability. We applied CSS and other previously published integration approaches to a complex cerebral organoid scRNA-seq dataset [11], where the data was affected by technical variation due to organoid, batch, and iPSC line/ human individual. Altogether the dataset contained scRNA-seq data from 20 two month old human cerebral organoids, each with a different cell type composition, from seven different ESC/iPSC lines in four batches of in total eleven experiments (Fig. 2a). UMAP embedding of the data without any integration method reveals segregation of cells due to cell type (e.g. brain region), batch, and line/individual (Fig. 2b). Previously, RSS with human fetal BrainSpan RNA-seq data as reference was used to integrate the data from different experimental batches [11], allowing interpretable cell type annotations (Supplementary Fig. 2, Fig. 2c). In this case, there are neuronal differentiation trajectories from multiple brain regions including cortex, GE, and non-telencephalic regions. This RSS-based cell type annotation, as provided in the original study [11], was considered as the primary reference annotation for a comparison of CSS with other integration methods. To avoid putative bias in our comparisons, we developed an alternative semi-automated integration-free strategy to annotate cerebral organoid scRNA-seq data and applied it to this data set, which resulted in the secondary reference annotation (Supplementary Fig. 2).

**Fig. 1.**
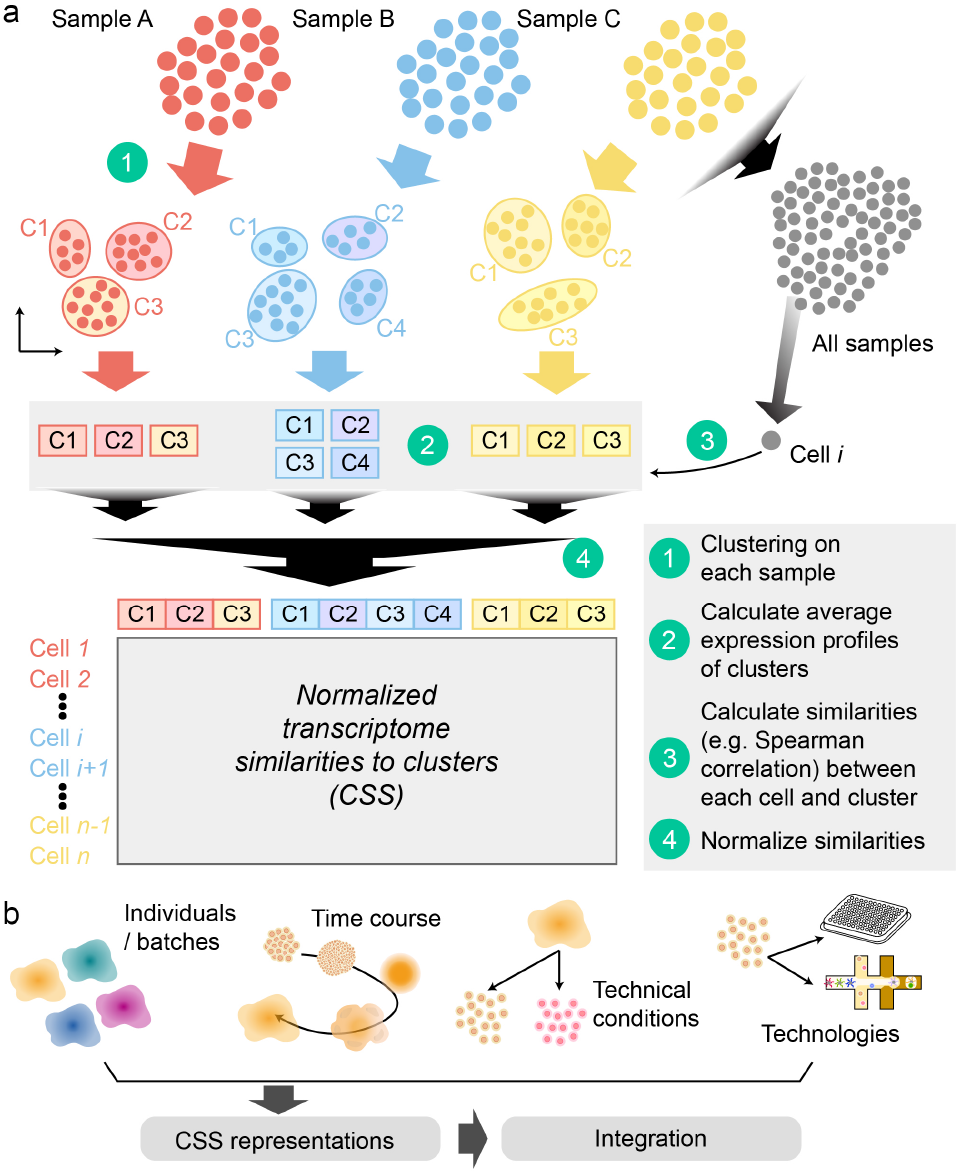
Schematic illustration of Cluster Similarity Spectrum data representation. (a) First, clustering is performed on each sample (A, B, C) separately. Second, the average expression profile of each cluster is calculated. Third, the correlation (e.g. Spearman or Pearson’s) is calculated between each single-cell transcriptome and the average transcriptome of each cluster to obtain a given cell’s transcriptome similarity. Fourth, for each cell the resulting similarities are normalized across clusters of each sample, and the normalized similarities to different samples are concatenated for the final cluster similarity spectrum (CSS) of the cell. The CSS vector representation of each cell is used for downstream clustering, embedding, projection, and other analyses. (b) We applied CSS to integrate human cerebral organoid data from different individuals, batches, experimental and technical conditions, and technologies.

**Fig. 2.**
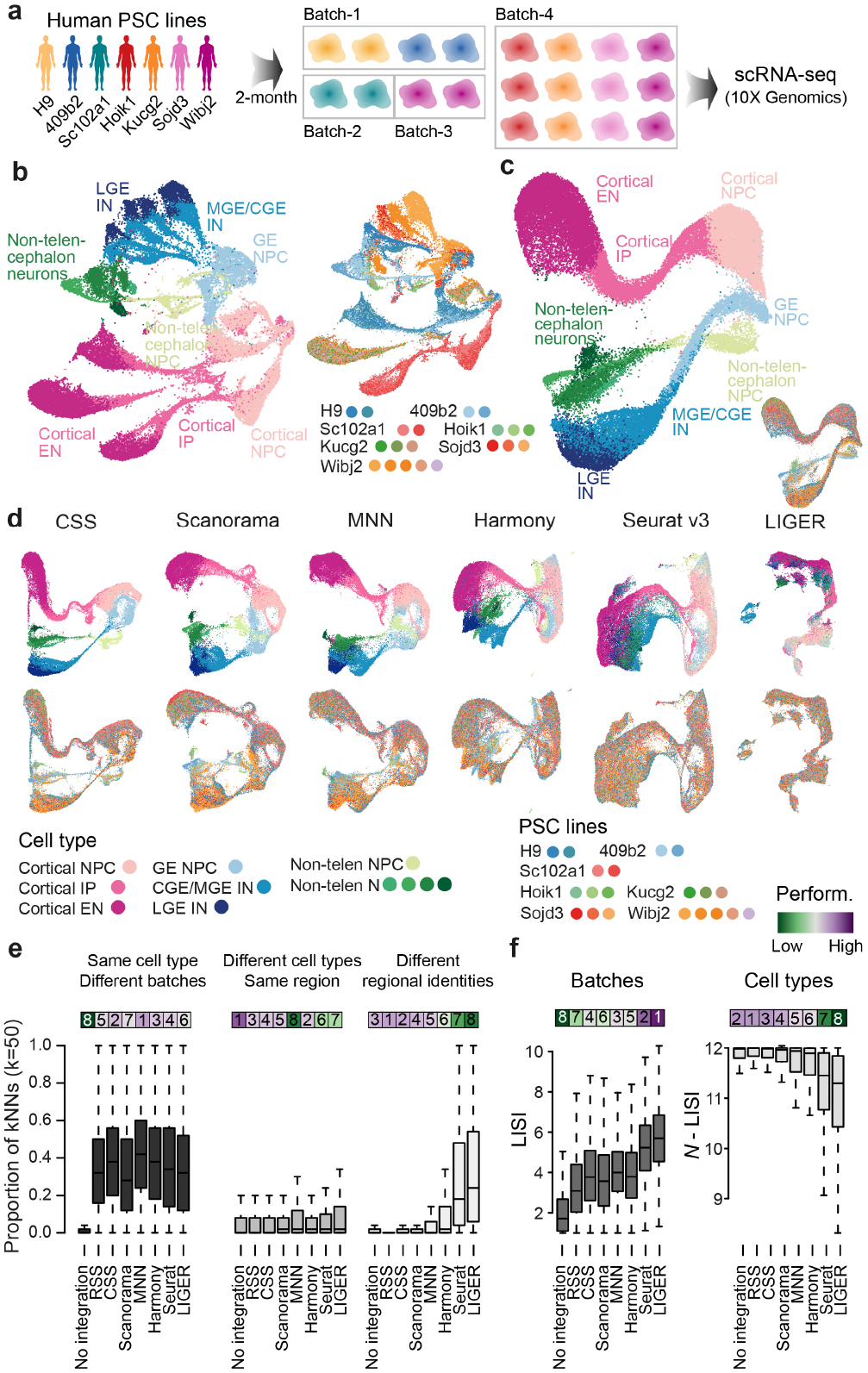
Comparison of CSS to different integration methods to resolve cell type heterogeneity in cerebral organoid scRNA-seq data. (a) Schematic of the experimental design. 20 two-month-old cerebral organoids were generated from seven different human PSC lines in four batches, and scRNA-seq data was generated using 10x Genomics. (b) UMAP embedding of cells without integration, with cells colored by RSS-based cell type annotation (left) and organoid of origin (right). (c) RSS with the fetal BrainSpan RNA-seq data as the reference integrates cells of different organoids, and is used for the comparison. The RSS-based UMAP embedding is colored by cell type annotation and organoid, respectively. (d) UMAP embeddings based on CSS and five other integration methods, colored by RSS-based cell type annotation (top row) and organoid (bottom row). (e) Proportions of cell neighbors in each annotation category across all cells in the data set. Cell neighbors are defined as the top 50 cells with the shortest Euclidean distances from the cell in PCA (no integration, left) or the seven different integrated representation spaces. Bars on top are colored by the median proportions, with the numbers showing ranks of different methods. (f) Distribution of LISI scores of organoid batches (left) and N-LISI scores of cell types (right) for all cells, without integration or with each of the seven integration. Bars on top are colored by averages across all cells, with the numbers showing ranks of different methods.

Next we applied CSS, MNN, Scanorama, Harmony, Seurat v3, and LIGER to this cerebral organoid dataset to compare the performance of different integration approaches. The UMAP embeddings of all four integration methods show that each method largely improves mixing of cells from different organoids and batches (Fig. 2d). For each cell in the data set, we calculated the proportions of neighboring cells being of the same cell type but from different batches. Here neighboring cells of one particular cell were defined as the 50 cells with the smallest Euclidean distances to the cell of interest in the integrated space. All six integration methods substantially increased the proportions, indicating that they all performed well in cell mixing (Fig. 2e). However, Seurat v3 and LIGER suffered from an over-correction problem, where cells of different differentiation stages or of different brain regions were grouped together (Fig. 2d-e). On the other hand, cell states remained well separated but nicely mixed between batches when the data was integrated with CSS, Scanorama, MNN and Harmony, providing results comparable to RSS (Fig. 2d-e). Consistent with this observation, we also computed Local Inverse Simpson’s Index (LISI) [8] of organoids and cell types with different integration methods and found similar results. (Fig. 2f). Among the four integration methods, CSS provided the best balance between mixing cells from different batches and separating different cell types, ranking the top or the second in all the performance metrics (Fig. 2e-f, Supplementary Fig. 2). All the observations were consistent regardless of which cell type annotation was used.

### CSS integrates time course data

ScRNA-seq technologies provide snapshots rather than longitudinal measurements of sample heterogeneity. To study dynamic processes, measuring multiple different samples representing different time points is often mandatory. In such a scenario, temporal variation is unavoidably confounded by differences between individual samples and experimental batches, and it becomes difficult to disentangle biological and technical portions of variation.

To assess whether CSS can integrate time course scRNA-seq data, we applied it to a dataset of cerebral organoid development from pluripotency [11] (Fig. 3a). This dataset includes seventeen samples from two PSC lines, covering seven different developmental stages. In comparison, Scanorama, MNN, Harmony, Seurat v3 and LIGER were also applied to the same data set. The UMAP embedding shows that without any integration cells of samples from different time points, particularly PSC, embryoid body (EB), neuroectoderm (NEcto) and neuroepithelium (NEpith) stages, separate from each other, forming distinct cell groups (Fig. 3b). Differences between the two PSC lines are also substantial.

**Fig. 3.**
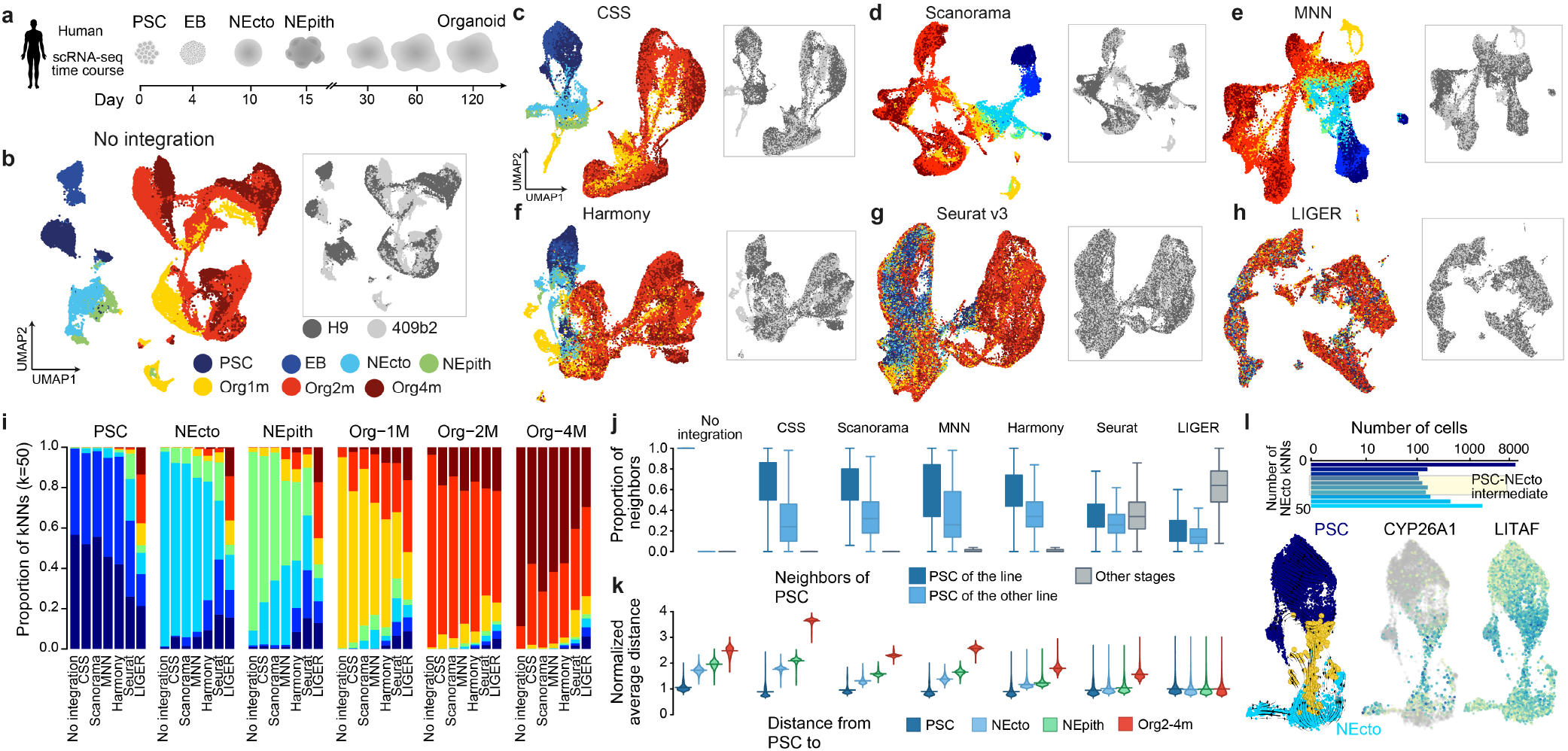
CSS integrates time-course scRNA-seq data of cerebral organoid development from pluripotent stem cells. (a) Schematic of the experimental design. PSC, pluripotent stem cell; EB, Embryoid Body; NEcto, Neuroectoderm; NEpith, Neuroepithelium. (b) Joint analysis of the time course cerebral organoid scRNA-seq data without integration. Dots, each representing one cell, are colored by sample time points (left) and PSC lines (right), respectively. Expression patterns of example markers are shown as feature plots. Org, Organoid; m, month. (c-h) Joint analysis of time course scRNA-seq data with (c) CSS, (d) Scanorama, (e) MNN, (f) Harmony, (g) Seurat v3 and (h) Harmony integration. The cells in the UMAP embedding are colored by sample time points (left) and PSC lines (right), respectively. (i) Stacked bar plots showing average proportions of cells from different time points in the neighborhood of cells in different time points when no integration or different integration methods are used. (j) Proportion of PSC neighbors in different annotation categories. PSC is the union of cells in PSC and EB time points. Neighbors are defined for each cell as top 50 cells with the shortest Euclidean distance in PCA (no integration) or other integrated representation spaces. (k) Average Euclidean distances from PSC to cells at later time points in PCA and different integrated representation spaces (bottom). (l) Intermediate cells between PSC and neuroectoderm (NEcto) stages are defined as those with comparable proportion of PSC and NEcto neighbors in the CSS space (top). RNA velocity analysis (bottom left) supports a potential cell state transition from PSC (dark blue) to NEcto (light blue) via the identified intermediate state (yellow cells). Expression patterns of two genes marking the intermediate state, CYP26A1 and LITAF, are shown (bottom right).

Strikingly, we find that CSS, Scanorama, MNN and Harmony largely integrate cells of the early time points including PSC to NEpith stages, as well as cells of different line origin, without disturbing later time points where different neuronal cell types have emerged (Fig. 3c-h). In comparison, Seurat v3 and LIGER encountered strong over-correction problems and largely mixed PSCs with NPCs in cerebral organoids (Fig. 3g-i).

To quantitatively compare different integration results, we focused on the PSC/EB time points, which are known to be distinct from cell states in the later time points [15, 16]. As a measure for comparison, we quantified the line and time point composition of cells neighboring each PSC/EB cell. All six integration methods intermixed cells from different PSC lines. However, a substantial proportion of cells of non-PSC/EB stages became neighboring cells of the PSC/EB population when Seurat v3 and LIGER were used (Fig. 3i-j), implying an over-correction problem. We also calculated average distances between each PSC/EB cell and cells at different time points (Fig. 3k). The distances to PSCs increased along the developmental time course for five out of the six tested integration methods with LIGER being the only exception. In general, CSS, Scanorama, MNN and Harmony substantially outperformed Seurat v3 and LIGER. We also extended the quantitative comparison to other cells by looking at time point and line labels of the 50 nearest neighbors of each cell in different integration spaces (Supplementary Fig. 3). All the integration methods substantially increased the enrichment of cell neighborhoods from different cell lines, while only CSS, Scanorama and MNN resulted in net gain by avoiding over-mixing cells from different time points (Supplementary Fig. 3). In addition, CSS and MNN are the two best methods for increasing connectivities between cells from nearby time points relative to cells from distant time points (Supplementary Fig. 3).

Interestingly, a continuous linkage was observed between earlier time points, e.g. from PSC/EB to NEcto cells in the UMAP embedding (Fig. 3c-h) when CSS, Scanorama, MNN or Harmony was used. We defined intermediate cells between PSC/EB and NEcto, as well as those between NEpith and one-month-old organoids, based on different integration methods, and compared their molecular signatures with the non-intermediate cells (Supplementary Fig. 3). We found that intermediate cells defined by CSS and Harmony presented the most unbiased distribution along the trajectory (Supplementary Fig. 3). We next focused on transcriptome signatures of intermediate cells between PSC/EB and NEcto defined by CSS representation. Those cells (in total 295 cells) may represent the transition state of neural induction. RNA velocity analysis using scVelo [17] also supported the transition potential from PSC to NEcto cells (Fig. 3l). Comparing those transition cells to both PSC/EB cells and NEcto cells resulted in differentially expressed genes such as CYP26A1 and LITAF (Fig. 3l). Among them, CYP26A1 encodes for a retinoic acid-metabolizing enzyme and has been shown to be essential for body patterning and brain development[18–20]. Knocking out this gene has been reported to reduce embryonic stem cell differentiation in response to retinoic acid [21].

We applied the above six integration methods including CSS, Scanorama, MNN, Harmony, Seurat v3 and LIGER to another time course scRNA-seq data set of human retinal organoids and primary retina, where cells were annotated as 11 different cell types [22] (Supplementary Fig. 4). The data set consists of 24 samples covering 20 different time points, including four retina organoids cultured for 24-59 days, 17 fetal retina samples aged between gastrulation week 9 to 27, one retina sample at postnatal day 8 and one adult retina. Similarly, all the integration methods substantially improved mixing between different samples, while CSS, Scanorama and MNN performed the best, preserving the temporal order of samples. These three methods also performed best by avoiding mixing cells of different cell types (Supplementary Fig. 4). We also investigated whether the integration methods retained the identity of intermediate cell states rather than mixing them with their progenitor or differentiated cell types, by focusing on two annotated intermediate precursor cell types (AC/HC precursors, BC/Photo precursors) (Supplementary Fig. 4). We found that CSS performed the best in remaining identities of intermediate cell states, as well as preserving its linkage with its biological progenitor and differentiated states.

These results together suggest that CSS is able to resolve a developmental intermediate state.

### CSS integrates data across experimental conditions

To determine if CSS can integrate data across experimental conditions, we generated scRNA-seq data from fresh and methanol-fixed cerebral organoid single-cell suspensions (Fig. 4a). Methanol fixation was applied to half of the dissociated cells prior to the scRNA-seq experiment. ScRNA-seq experiments were performed on the fixed and unfixed cells separately. The data suggested moderate differences between the two experimental conditions (Fig. 4b). We applied RSS using the fetal BrainSpan RNA-seq data as reference to integrate both conditions, define cell clusters and annotate cell populations (Fig. 4c). In addition, we also applied the integration-free cell type annotation strategy described above to get the alternative cell type annotation that was not reliant on RSS.

**Fig. 4.**
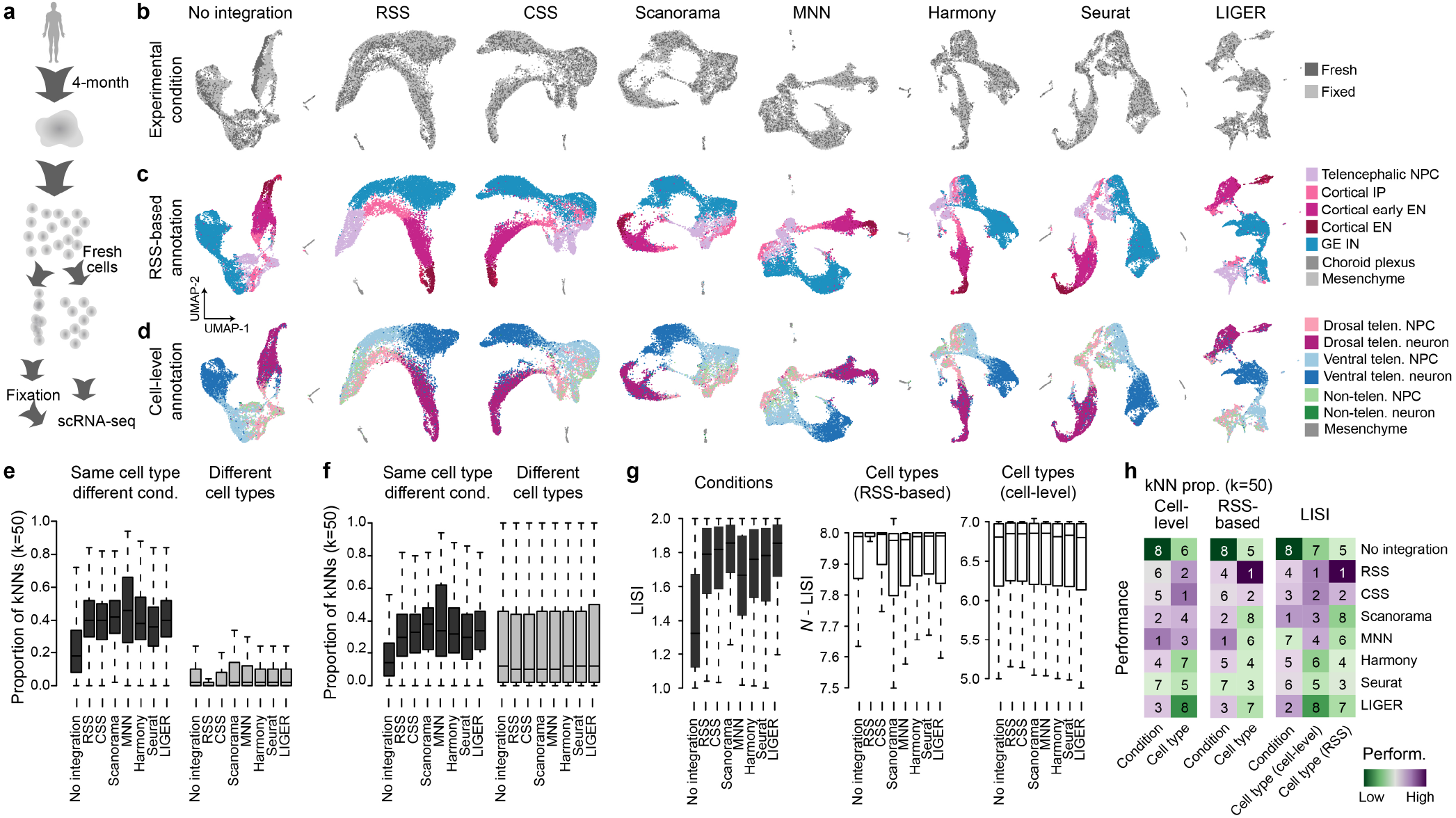
Integration of scRNA-seq data of cerebral organoids with fixed and unfixed experimental conditions. (a) Schematic of the experimental design. Fresh and methanol-fixed single-cell suspensions of the same four-month-old cerebral organoid were measured using scRNA-seq. (b-d) UMAP embedding of the scRNA-seq data before and after integration with the seven integration methods. Cells are colored by (b) experimental conditions, (c) RSS-based cell type annotations, and (d) model-based predicted cell type annotations. (e-f) Proportion of neighbors of each cell in different categories, with neighbors defined as 50 cells with the shortest Euclidean distances with a cell in PCA (no integration) or different integration spaces. Cell types are annotated based on (e) clustering on RSS or (f) model-based prediction. (g) LISI-based scores of conditions (left), RSS-based cell type annotation (middle) and model-based cell type annotation (right) with no or different integration methods. (h) Heatmaps showing summarized performance metrics. Numbers show the ranking of different integration methods based on different metrics.

We then applied CSS, Scanorama, MNN, Harmony, Seurat v3 and LIGER and estimated performances of the methods. We compared the cell type and experiment condition of each cell with its 50 nearest neighbors defined using different integration methods. LISI was also computed for experiment condition and cell type annotation to compare different integration methods. The results suggest that all six methods significantly increased the mixing of cells from different experimental conditions (Fig. 4e-h). Over-integration remained low in general, but it is worth mentioning that CSS outperformed other methods to avoid mixing of different cell types (Fig. 4h). This result indicates that in this common experimental scenario where fixation is performed, CSS performs similarly well as the other integration methods.

### CSS integrates scRNA-seq data generated by different technologies

We next sought to assess the performance of CSS for the integration of scRNA-seq data generated by different technologies. We compared cerebral organoid scRNA-seq data generated using 10x Genomics (one cerebral organoid, 4512 cells), Fluidigm C1 and Smart-Seq2 (C1/SS2, 685 cells) [11], or inDrops (5847 cells, generated in this study). Analysis on the three data sets separately revealed their cell composition, allowing annotation of cells into nine different groups, including NPCs and neurons of the dorsal telencephalon, the ventral telencephalon and of non-telencephalic regions, choroid plexus, mesenchymal-like cells and epithelial-like cells (Fig. 5a). Joint analysis without integration showed that single cell transcriptomes primarily segregate out based on the underlying scRNA-seq technology.

**Fig. 5.**
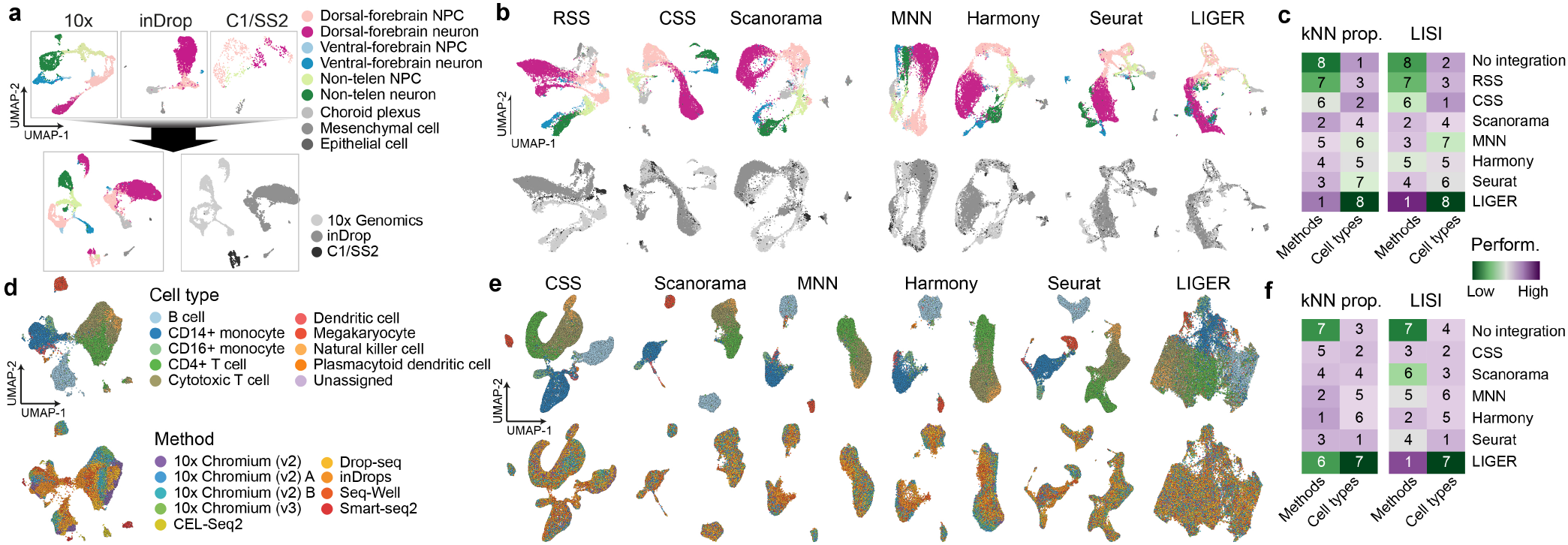
CSS integrates scRNA-seq data generated using different technologies. (a) Joint analysis of cerebral organoid scRNA-seq data generated using different technologies without integration. Cell type annotation is performed in each data set separately (top). Joint UMAP embedding is generated without integration, and colored by cell types (bottom left) and technologies (bottom right), respectively. (b) UMAP embeddings based on different integration methods, colored by cell type annotation (top) and technology (bottom). (c) Heatmaps showing summarized performance metrics. Numbers show the ranking of different integration methods based on each metric. (d) Joint analysis of public PBMC scRNA-seq data generated using different methods without integration. UMAP embeddings are colored by annotated cell types (top) and methods (bottom). (e) UMAP embeddings based on different integration methods, colored by cell type annotation (top) and methods (bottom). (f) Heatmaps showing summarized performance metrics. Numbers show the ranking of different integration methods based on different metrics.

We integrated the three data sets using RSS (fetal BrainSpan RNA-seq data as reference), CSS, Scanorama, MNN, Harmony, Seurat v3 or LIGER. All methods, except RSS, largely improve mixing of cells from different data sets/technologies (Fig. 5b-c), although over-integration was apparent in the case of LIGER integration (Fig. 5c, Supplementary Fig. 5). Scanorama, MNN, Harmony and Seurat v3 performed comparably well in terms of mixing cells from different data sets and better than CSS. On the other hand, CSS performed the best with regard to balancing improvements in mixing cells from different technologies and retaining clear cell type heterogeneity (Fig. 5c, Supplementary Fig. 5).

All the above analyses were performed on cerebral organoid scRNA-seq data sets, which represent developmental processes with continuous molecular changes and trajectories. Next, we sought to estimate how CSS performed on data sets with more distinct cell types. We downloaded the public scRNA-seq data of human PBMCs [23], and applied CSS, Scanorama, MNN, Harmony, Seurat v3 and LIGER to integrate cells measured by different methods (Fig. 5d-e). Over-integration occurred when LIGER was used (Fig. 5e-f, Supplementary Fig. 5), while all the other methods including CSS largely improved mixing of similar cells measured by different methods (Fig. 5f, Supplementary Fig. 5). CSS showed good performance in avoiding mixing of different cell types, ranking second. In general, Seurat v3 performed the best, but other methods including CSS, except for LIGER, were not far behind based on these metrics.

### CSS allows query data projection to the reference scRNA-seq atlas

As most of the integration methods require data specific transformation of the expression matrices, any integration procedure needs to be rerun when a new data set is introduced. In contrast, CSS representation relies on normalized expression levels without additional data transformation and therefore allows for the projection of new query data to a CSS-represented reference atlas. We tested the feasibility of such a procedure with the scRNA-seq data set of two-month-old cerebral organoids introduced in Figure 1. In brief, we first extracted cells of two organoids from the Sc102a1 iPSC line as the query data. Next, we built a CSS-integrated reference atlas using the remaining scRNA-seq data (Fig. 6a). The corresponding CSS representations of the query cells in the Sc102a1 organoids were then calculated. The projected UMAP embedding of the query cells, as well as their projected cell type labels, were obtained with the intrinsic UMAP projection mechanism and a k-nearest-neighbor (kNN, k=50, defined in the CSS-space) classifier, respectively. The projected UMAP embedding is consistent with the previous analysis showing that the two Sc102a1 cerebral organoids mostly consist of cortical cells (Fig. 6b). More importantly, using the kNN classifier to predict the cell type of the query cells based on the reference cell annotation resulted in the right prediction for 89.4% of the query cells. 8.9% of query cells were predicted to be a different cell state than previously annotated, but along the right fate trajectory, i.e. cortex, GE or non-telencephalon (Fig. 6b). This result suggests that the CSS-based data projection procedure is technically straightforward and reliable.

**Fig. 6.**
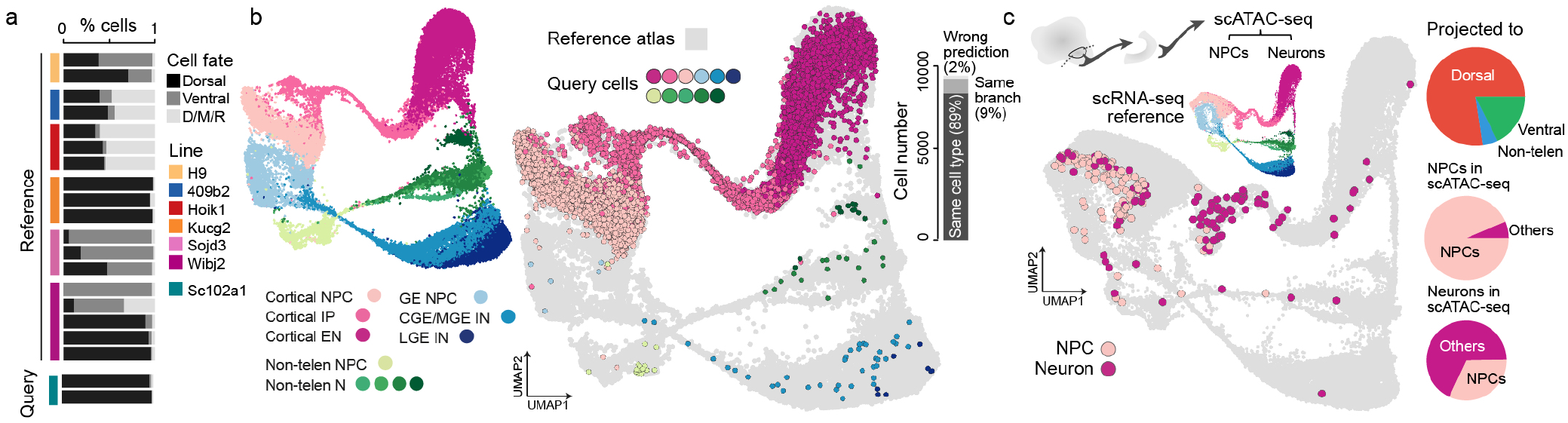
Projection of scRNA-seq and scATAC-seq data to reference scRNA-seq atlas using CSS. (a) Projection of scRNA-seq data of two Sc102a1 cerebral organoids to the reference atlas built using scRNA-seq data of the other eighteen cerebral organoids. The barplot shows the proportion of cells per organoid that are located on the dorsal or ventral telencephalon or diencephalon, mesencephalon and rhombencephalon (D/M/R) neuronal trajectory. (b) UMAP embedding of the reference atlas based on CSS representation. Cells are colored by cell type annotation (left). Cells in the Sc102a1 organoids are projected to the reference atlas with cells in the projected embedding colored by their previous (see Figure 2) cell type annotation (middle). Proportion of cells for which the predicted cell type is consistent or inconsistent with the previous annotation is shown in the barplot (right). (c) Projection of scATAC-seq data of dissected cortex-like areas in cerebral organoids to the scRNA-seq reference atlas including twenty cerebral organoids. Projected loci of cells in the scATAC-seq data set is shown in the reference UMAP embedding, colored by the independent cell type annotation. Pie charts show the proportion of cells in the scATAC-seq data set projected to dorsal, ventral or D/M/R neuronal branches (top right), the cell type composition of cells predicted as NPCs (middle right), and the cell type composition of cells predicted as neurons (bottom right).

We further sought to determine whether the projection procedure can be applied to scATAC-seq data as query data. We used scATAC-seq data (Fluidigm C1) of micro-dissected cortex-like structures in human cerebral organoids [11] (Fig. 6c). For each cell, its chromatin accessibility pattern was summarized into a gene activity profile, defined as the enrichment of accessible regions in the promoter and gene body of different genes. CSS representations of those cells towards the scRNA-seq reference atlas were then calculated based on their gene activity profiles. The projected UMAP embedding and cell type labels were then compared to the annotation based on the scATAC-seq data (Fig. 6c). We found that most of the cells (77.4%) in the scATAC-seq data were projected to the dorsal telencephalic branch which matches with the experimental design. Using the cell type annotation by clustering of the scATAC-seq data as the benchmark, most of the cells in the scATAC-seq data that annotated as NPC (93.5%) projected to NPC in the reference atlas, while the majority of the cells annotated as neurons projected to IP or neurons in the reference (67.8%). These results show that CSS-based representation and projection of scATAC-seq to a scRNA-seq reference atlas is therefore possible and can be informative for cell type annotation and interpretation of the scATAC-seq data.

## Discussion

Here we present CSS, a simple but powerful data transformation strategy which can be used to integrate multiple data sets based on the comparison to intrinsic references. It represents cells by their transcriptomic similarity to cell clusters in different samples, with similarities to clusters within the same sample normalized. This representation can largely eliminate the influence of random technical variation across different samples. CSS is not the first method to propose usage of correlation to represent cell signatures. Other methods, e.g. scBatch [24] and RCA [12] have used the pairwise correlation matrix between cells to assist data integration. However, calculation of pairwise correlation greatly affects the scalability of these methods, making them difficult to be applied to large scRNA-seq data sets with tens of thousands or even more cells. This is not a problem in CSS, as cells are correlated to transcriptomic profiles of a limited number of clusters instead of every other cell in the data, allowing it to be applied to large scale scRNA-seq data.

We applied CSS to reanalyze complex scRNA-seq data of developing human cerebral organoids and a public PBMC scRNA-seq data set, as well as the newly generated paired cerebral organoid scRNA-seq data with fixed and unfixed experimental conditions. In our assessment, CSS successfully integrated scRNA-seq data of different experimental/technical conditions, cell lines, experimental batches and technologies, while retaining cell type heterogeneity. Compared to the commonly used tools Scaronama, MNN, Harmony, Seurat and LIGER, CSS performed similarly well or outperformed the other integration methods. In our benchmark, it provided comparable performances as the other methods in batch effect correction, while showing the best performance in retaining cell type heterogeneity especially in the cerebral organoid data sets representing data with continuous trajectories and intermediate states.

In principle, CSS representation corrects for random variation, which does not influence relative similarities of different cell types, across samples. In other words, any variation which changes the patterns of transcriptomic similarity, including samples at different conditions, are likely to remain. This behavior helps, in certain scenarios, e.g. when integrating time course samples from different temporal stages, and creates an integrated embedding allowing visualization of any variation on cells which affects their similarities to cell types in different samples.

On the other hand, the same feature could become a limitation of CSS. For instance, CSS may fail to integrate data across conditions or species, as there are likely differences among cells in those samples which affect their similarity spectrums. In that case, other integration methods which maximize correlations among different samples or conditions, e.g. Seurat and LIGER, may be the preferred solution. In addition, as CSS introduces a normalization across clusters of each sample, it requires at least moderate heterogeneity of the integrated samples so that the normalized spectrum is meaningful. When such a condition is not met, other methods may provide better results.

One useful feature of CSS representation is its ease of applicability to new data, which makes projections of new data to the CSS-represented reference data feasible. In this study, by splitting the human cerebral organoid scRNA-seq data into reference and query data and applying CSS for data representation, we showed that the CSS projection of query data is reliable and accurate, in terms of both the projected UMAP embedding and the transferred cell type labels. In addition, our work on projecting human cerebral organoid scATAC-seq data to the corresponding scRNA-seq reference further suggests that CSS can help linking different single-cell genomic data modalities and assist with annotation across modalities. Altogether, these features highlight the utility of CSS as a simple yet powerful approach for integration of complex single-cell sequencing data sets.

## AUTHOR CONTRIBUTIONS

ZH implemented the method and performed the analysis. AB performed scRNA-seq experiments using inDrop. SE performed organoid fixation and the associated scRNA-seq experiment. ZH, JGC, BT designed the study and wrote the manuscript.

## ACKNOWLEDGEMENTS

We thank E. H. Gustafson, S. Wolfinger and J. A. Knoblich of IMBA, Vienna for providing the cerebral organoids for the inDrops experiment. We thank L. Mazutis, J. Nainys, D. Kučiauskas, K. Simutis from Vilnius University for assisting with in-Drops platform setup and for providing barcoded hydrogels (Droplet Genomics), J. Kageyama and M. Dannemann for the computational support, and S. Jansen and S. Kanton for experimental support. This project has been made possible in part by the Chan Zuckerberg Initiative DAF (grant CZF2017-173814), an advised fund of Silicon Valley Community Foundation, European Research Council (Anthropoid-803441, J.G.C.; Organomics-758877, B.T.), Swiss National Science Foundation (Project Grant-310030_184795, J.G.C).

## DATA AVAILABILITY

The single-cell RNA-seq data using inDrop and the single-cell RNA-seq data of the fixation experiment were deposited on ArrayExpress with the accession number XXXXX, and Mendeley Data with DOI: 10.17632/3kthhpw2pd.1.

## Methods

### Principles of RSS and CSS

In general, the calculation of RSS and CSS includes the following steps (Supplementary Fig. 1). For RSS calculation, highly variable genes are first identified in the reference data. Reference samples are represented by their average expression profiles of the identified genes. Next, Pearson correlation coefficients are calculated, between each cell and every reference sample. Finally, resulting correlations are normalized across reference samples for each cell.

For CSS calculation, cells from different samples are first separated and clustered. In this study, we used the same Louvain graph-based clustering method as used in Seurat. In brief, a k-nearest neighbor (kNN) graph, weighted by Jaccard index, was firstly constructed for cells from each sample. In the results presented in this study, highly variable gene identification (using the vst method in Seurat), but other methods can also be used to generate the gene list for CSS calculation, e.g. the union set of the most correlated genes to the top PCs, or the consensus variable gene list of different samples. Principal component analysis (PCA) was then done before cells are subset into each sample, but doing PCA on each sample separately is also supported. The implementation of Louvain clustering in Seurat was then applied to each graph to get cell clusters of different samples. The clustering resolution influences the clustering result and therefore affects the CSS calculation, but the effect is not substantial as the resulting kNN neighborhoods are very robust to this parameter (Supplementary Fig. 2). The average highly variable gene expression profile of each cluster in each sample is calculated. For each cell, its similarity to each of the clusters in all samples is then calculated. In principle, any similarity metric can be used. Spearman and Pearson correlation coefficients between two transcriptome profiles are the two similarity metrics that are currently supported natively. In this study, we used Spearman correlation coefficient for the scRNA-seq data of cerebral organoids for its robustness to outliers, and Pearson correlation coefficient for scRNA-seq data set of PBMC to maximize the power to distinguish distinct cell type signatures (Supplementary Fig. 5).

Afterwards, a normalization procedure is applied to neutralize global differences of similarities to clusters of different samples. Two different strategies are proposed here. One option is z-normalization (i.e. centralize the data at zero and scale by the standard deviation) to the cell’s similarities to clusters of each sample. This method was used in all the scRNA-seq data of cerebral organoids in this study. We also propose the second option of correlation-based probability. First, the correlation kernel matrix [25] is calculated as:

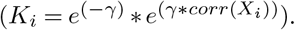

Here, *K_i_* is the resulting *N* ∗ *M_i_* kernel matrix, where *N* is the total number of cells and *M_i_* is the number of clusters in sample *i*. *γ* defines the sharpness of the normalized probabilities. A large *γ* amplifies the differences of similarities to different clusters. It is then transformed into a probability matrix by normalizing each row to have a sum of one. This method was used in the PBMC scRNA-seq data in this study. As the last step, for each cell, its normalized similarity spectrums to different samples are then concatenated as the final CSS representation, which is a *N* ∗ *M* matrix where N is the total number of cells and *M* is the total number of clusters in all samples.

The dimensions of CSS increase along with the number of samples to integrate, which affects the scalability of using CSS for following analysis. In addition, some technical variance among samples to be integrated may introduce bias towards certain cell types and fail to eliminate during the normalization procedure. To further deal with these two issues, an optional PCA can be applied to the CSS representation matrix to further reduce the data dimension and condense the information. This step was used in the PBMC scRNA-seq data in this study, with the first 10 PCs being used for constructing the UMAP embeddings and kNN network (Supplementary Fig. 5).

The underlying hypothesis of both RSS and CSS is that the undesired confounding factors, e.g. read coverage of different cells, introduce random perturbation to the observed transcriptomic measures which are not correlated with cell type or cell state identities. Cells with the same identity therefore share similar spectrums, after normalization across similarities to different references. In other words, the normalized similarity spectrum minimizes the global differences across cells introduced by the random perturbation. Additionally, as CSS considers all clusters in different samples as references, normalization is done separately across clusters of different samples, so that global differences across samples can be largely eliminated.

### Single-cell RNA-seq data generation of cerebral organoids in different experimental conditions

We acquired the human induced pluripotent stem cell (hiPSC) lines Kucg2 and Sojd3 from the HipSci resource, cultured and differentiated them to cerebral organoids following the same protocol as reported [11]. One organoid from each line at 116-day-old was dissociated as described previously [11]. Cell suspension of each organoid was split into two aliquots. Cells in one aliquot per line were pooled, together with cells of another human cerebral organoid of line Wibj2, also from HipSci resource, and one chimpanzee cerebral organoid, and diluted for an appropriate concentration to obtain approximately 10000 cells in one lane of a 10x microfluidic chip device. Methanol fixation was applied to cells in the remaining aliquots, following the same procedure as described previously [20]. Cells were kept at 4°C all the time. Briefly, between 1 and 2 × 106 cells from a filtered single-cell suspension of cerebral organoids, were pelleted at 300 x g for 5 min. The cell pellet was resuspended in 200 μl HBSS without calcium and magnesium (Gibco). To avoid clumping, 800 ul of methanol (pre-chilled to −20 °C) was added to the cells dropwise while gently vortexing the cell suspension at lowest speed. The methanol-fixed cells were kept on ice for 15 min and then stored at −80 °C for seven days. Afterwards, the fixed cells were rehydrated following the procedure described [20]. In brief, cells were moved from −80 °C to 4 °C and kept on ice during the procedure. Cells were then pelleted at 3000 × g, resuspended in DPBS with 2% (w/v) BSA and 1 U/μl RiboLock RNase Inhibitor (Thermo Fisher). Centrifugation and resuspending were repeated once more and cells were then passed through a 30 μm pre-separation filter (Miltenyi Biotec), counted and diluted to aim for 10000 cells in one lane of the microfluidic device (ChromiumTM Single Cell 3’ Solution v2, 10x Genomics). Libraries were sequenced on Illumina’s Hiseq2500 platform in paired-end mode (26+8 bp, 100 bp).

### Single-cell RNA-seq data generation using inDrop

Cerebral organoids were generated from an edited version of the embryonic stem cell line H9, following the protocol previously described [21]. Using CRISPR editing, double-stranded breaks were introduced in the first exon of the PLCB1 gene. One allele contained a 4bp deletion, the other allele a 19 bp insertion. Both mutations led to frame shifts and premature stops. At day-150 since culturing, two cerebral organoids were dissociated into single-cells using a papain-based neural tissue dissociation kit (Miltenyi) as described in Miltenyi’s Neural Tissue Dissociation protocol. To increase the accessible surface for the dissociation enzyme organoids were cut into pieces and washed up to 3 times in 1x HBSS without Ca^2+^ and Mg^2+^ (HBSS w/o, Sigma). In order to get sufficient disaggregation samples were gently triturated using wide-bore pipette tips and p1,000 and p200 pipettes. Next, a single-cell suspension was generated by removing clumps of cells filtering suspension through 30 and 20 μm diameter strainers. The single-cell suspension was washed up to 3 times in HBSS w/o. Samples were centrifuged at 300g for 5 min and resuspended in HBSS w/o. Finally, cell viability was determined with Trypan blue solution (0.4%) using an automatic cell counter (Countess, Invitrogen). For the single-cell RNA-seq experiment cells were diluted to 90,000 cells/ml in 15% Optiprep and 0.02% BSA in PBS.

Single-cell transcriptome barcoding was performed using inDrops [3] and following the protocol by Zilionis et al, 2017 [22]. Shortly, generated single-cell suspension was co-encapsulated with RT-lysis mix and barcoded hydrogel beads (Droplet Genomics) into 3-4 nl droplets. After cDNA synthesis was performed, droplets were broken and cDNA libraries were prepared by second strand synthesis, linear amplification by in vitro transcription, amplified RNA fragmentation, reverse transcription and PCR. Prepared cDNA libraries were sequenced paired-end (100 bp, 50 bp) on an Illumina HiSeq2500 platform on 2 lanes.

### Data retrieve and processing

ScRNA-seq data sets of human two-month-old cerebral organoids and cerebral organoid developmental time course from PSC, as well as the metadata and cell type annotation, were retrieved from ArrayExpress (accession E-MTAB-7552) [11] The scATAC-seq data was retrieved from ArrayExpress (accession E-MTAB-8089) [11]. Quality control was done with the same procedure as described in the original publication.

The scRNA-seq data set of retinal organoids and primary retina samples, including the metadata and cell type annotation, were retrieved from GEO (accession GSE138002) [22]. The retinal organoid sample cultured for 24 days was excluded from the analysis as it only contains 24 cells.

For the newly generated scRNA-seq data of fixed/unfixed experimental conditions, Cell Ranger, the set of analysis pipelines suggested by 10x Genomics, was used to demultiplex raw base call files of libraries by 10x Genomics to FASTQ files and align reads to the human genome and transcriptome (hg38, provided by 10x Genomics) with the default alignment parameters. Demultiplexing of human and chimpanzee cells in the unfixed sample was done based on genomic loci with diverged bases between human and chimpanzee, following the same procedure as described [11]. Only human cells were used in the later analysis. Demultiplexing of the three human lines in the unfixed sample was done using demuxlet [23], based on the genotyping information of lines downloaded from HipSci websites. Cells with the best singlet prediction being Wibj2 with likelihood no less than 10 higher than the second best singlet likelihood were discarded.

For the newly generated inDrops scRNA-seq data, preprocessing was done following the Drop-seq tools procedure. In brief, quality control to the FASTQ files was done by removing reads with multiple low-quality bases at cell or molecular barcodes, and polyA sequences (with at least six consecutive As) trimmed using Drop-seq tools (v1.12). Remaining reads were mapped to the hg19 human genome using STAR with default parameters. Count matrices were made using Drop-seq tools. Cells with less than 10000 reads were dropped from the analysis.

For the PBMC scRNA-seq data, we retrieved the Seurat object directly using the SeuratData package in R to download the data set “pbmcsca”.

For all the data sets in this study, normalization (log normalization with default parameters) and highly variable feature identification (vst method) was done by Seurat (v 3.1). After integration, the k-nearest neighbors (k=50) were identified using the nn2 function in the RANN R package with default parameters, which searches for cells with the smallest Euclidean distances with the given integrated data for each cell.

To get an automated cell type annotation of cells in the two-month-old cerebral organoids, we firstly applied the louvain clustering (implemented in Seurat, resolution = 0.6) to the first 20 PCs of the data set without integration, resulting in 25 clusters. Based on regional identity markers including *FOXG1*, *EMX1* and *DLX2*, prior regional identities of dorsal telencephalon, ventral telencephalon and non-telencephalon were assigned to cells in six, five and five clusters, respectively (Supplementary Fig. 2). 1000 cells from each identity were randomly selected, and a LASSO regression model (with multinomial family) was fitted to predict regional identity of a cell given its expression values of the highly variable genes. The trained model was applied to each cell in the data set for the predicted regional identity. Next, we calculated a neural progenitor (NPC) score, defined as the average expression level of 47 genes with higher expression level in NPCs than in neurons, and a neuron score, defined as the average expression level of 29 genes with higher expression level in neurons, for each cell. A cell with higher NPC score than neuron score was considered as an NPC, and a cell with higher neuron score was considered as a neuron (Supplementary Fig. 2). For each cell, the combination of its predicted regional identity and estimated NPC/neuron identity was used as its alternative cell type annotation. The same method was also applied to the scRNA-seq data set of cerebral organoids in different experimental conditions.

### Data integration

To integrate the human two-month-old cerebral organoid scRNA-seq data, RSS to the human fetal BrainSpan RNA-seq reference data was calculated as described in the original publication [11]. For CSS calculation, principal component analysis (PCA) was firstly applied to the data considering the top 5000 highly variable genes. CSS was then calculated, with different organoids as different samples for louvain clustering (with resolution of 0.6) implemented in Seurat, which took the top-20 calculated principal components (PCs) as the input. Scanorama was applied to the highly variable genes expression matrix with default parameters. MNN was applied to the highly variable gene expression with default parameters using the RunFastMNN wrapper function in the SeuratWrappers R package. Harmony was applied with the default parameters and the top-50 PCs, calculated in the same way as above, using the RunHarmony function in the harmony R package, to integrate different organoids. Seurat v3 was applied following its standard workflow of integration, using 5000 features for anchoring and top-30 PCs in the weighting procedure. LIGER was applied following basic commands tutorial, with variance threshold being 0.3, inner dimension of factorization being 20, convergence threshold being 5E-5, three restarts of integrative non-negative matrix factorization and clustering resolution of 0.4. The same parameters were also applied to the integration of the developmental time course scRNA-seq data of human cerebral organoids from PSC, and the integration of scRNA-seq data of human cerebral organoids in different experimental conditions (fixed/fresh). The only exception is the variance threshold in LIGER, which was set as 0.1 for the developmental time course data, and 0.01 for the two experimental conditions data. Such difference was made to keep the number of variable features used in LIGER integration similar.

To integrate the human retinal organoid and primary retina scRNA-seq data, the same procedure as above was used. The only exception is for LIGER calculation, we used the RunOptimizeALS and RunQuantileAlignSNF functions in the SeuratWrappers R package using the same setting as in the online vignette (https://htmlpreview.github.io/?https://github.com/satijalab/seurat.wrappers/blob/master/docs/liger.html).

To integrate the three scRNA-seq data sets generated by different technologies, 5000 highly variable genes were determined for each data set separately. Genes defined as highly variable in at least two data sets after excluding genes reported as cell-cycle-related genes were used for integration, accounted for 2984 genes, were used for integration. The variance thresholds in LIGER were 0.3, 1 and 0.8 for the 10x, inDrop and C1/SS2 data sets, respectively. Other parameters are the same as described above.

To integrate scRNA-seq data of different methods in the PBMC data set, 3000 highly variable genes were firstly identified. Similar procedures as above were used, except for CSS and LIGER. For CSS calculation, a clustering resolution of 0.4 was used for Louvain clustering of each sample, and Pearson correlation was calculated between each cell and each cluster. Similarity normalization was done via kernel probability transformation (γ=50). For LIGER calculation, we used the RunOptimizeALS and RunQuantileAlignSNF functions in the SeuratWrappers R package using the same setting as in the online vignette (https://htmlpreview.github.io/?https://github.com/satijalab/seurat.wrappers/blob/master/docs/liger.html).

### Quantitative metrics of integration performance

To quantify the performances of different integration methods on the human two-month-old cerebral organoid scRNA-seq data, we calculated k-nearest neighbors (kNN, k=50) for each cell in different integration space (RSS, CSS, Scanorama, MNN, Harmony-integrated top-50 PCs, Seurat-integrated top-20 PCs, and LIGER-based quantile aligned factor loadings). Based on the cell type annotation retrieved from the original publication, which is based on RSS, we counted the proportions of neighbors for each cell, which are annotated as the same cell type but of an organoid in a different experimental batch, a different cell type but with the same regional identity, and a different regional identity. A good integration method should increase the proportion of neighbors annotated as the same cell type but from a different batch, while keeping the proportion of neighbors annotated as a different cell type low.

In addition, Local Inverse Simpson’s Index (LISI) [8] was computed to access organoid mixing and cell type separation. LISI is used to determine the number of cells that can be drawn from a neighbor list before one batch is observed twice. The LISI scores range from 1 to *N*, where *N* is the total number of organoids or cell types in the dataset. In this study, we used LISI of organoids as a metric of batch correction, and *N* -LISI (*N* is the total number of cell types) as a metric of cell type separation. A fixed perplexity of 30 was used. For both metrics, a higher score indicates better performance.

For the developmental time course scRNA-seq data set of human cerebral organoids from PSC, kNN-based metrics similar as above was used but focusing on PSC (including PSC and EB time points), as it is the most distinct cell type from the others. The proportions of nearest neighbors of each PSC which was PSC of the same line, PSC of the other line, or cells from any other sample were calculated. A good integration method should show a high proportion of PSC neighbors being PSC of the other line and low proportion of cells from other samples. In addition, average distances between each PSC and cells at PSC/EB, neuroectoderm, neuroepithelium, and cerebral organoids at age of two-to-four months were calculated on each integration space. The resulting average distances were normalized to the mean of average distances between different cells at PSC stage to allow comparison of different integration methods.

To extend the assessment to not only the PSCs, but also other cells in the data set, we calculated the log-transformed odds ratio between the proportion of its kNN (k=50) of the same line and the proportion of this line in the whole data set (lOR), based on different integration as well as no integration. A higher lOR indicates better mixing of cells from different lines. A similar log-transformed odds ratio for each cell was calculated also for time points (tOR). The difference between lOR and tOR represents the excess of line mixing relative to time point mixing, which is expected to be positive in this data set, as both lines cover all the time points. In addition, for each cell we also calculated the log-transformed odds ratio of neighbor cells from the nearby time points (ntOR) and those from the distal time points (dtOR). A higher ntOR indicates better mixing of cells from nearby time points, while a more positive ntOR-dtOR suggests more proper temporal ordering.

For the developmental time course scRNA-seq data set of human retinal organoids and primary retina, kNN-based metrics similar as above was used. We counted the proportions of neighbors for each cell which were from the nearby time points. A higher proportion suggests improved data continuity. We also counted the proportions of neighbors from the nearby time points among neighbors from all the different time points. A higher proportion suggests better recovery of the proper developmental temporal order. The proportions of neighbors annotated as different cell types were also counted, which should remain low if the integration successfully avoids over-correction.

Similar as above, we calculated kNN for cells in different integration spaces to access the performances of different integration methods on different experimental conditions and technologies. Proportions of neighbors of each cell annotated as the same cell type but represent different experimental conditions or technologies, and cells annotated as a different cell type were calculated. As the alternative metrics, LISI scores of experimental conditions or technologies, and *N* -LISI scores of cell types, were also calculated.

In all the analysis, integration methods were ranked based on the average scores of metrics across all cells in the data set.

### Characterization of intermediate cell states in the time course cerebral organoid data

To identify intermediate cells between cells from different time points (e.g. PSC and neuroectoderm, neuroepithelium and one-month organoid), we firstly identified kNNs (k = 50) on the integrated space for each cell in the two time points of interest. Cells with at least 15 neighbors from each of the two time points are defined as cells at the intermediate state. This procedure was applied separately to PSC and neuroectoderm cells, as well as neuroepithelium and one-month-old organoid cells for their intermediate cells.

To benchmark the unbiasedness of intermediate cells identified based on different integration methods, we united cells identified with at least two integration methods, and looked for their kNNs (k=50) in different integration spaces. Non-intermediate cells at the two time points of interest which appeared for at least 10 times as neighbors of the intermediate cell union were used as the control cells. Differential expression (DE) analysis was done using the presto R package between control cells of the two time points. A signature score of each time point was calculated for each control cell and intermediate cells, as the average expression level of genes with significantly higher expression in control cells of the time point (AUC>0.7, average fold change > 1.2, detection rate difference > 30%, BH-adjusted two-sided Wilcoxon rank sum test P<0.01). The difference between the two scores was defined as the time point signature bias. Different integration methods were thus ranked based on the median time point signature bias of all the intermediate cells identified based on each integration method.

To further characterize molecular signatures of intermediate cells between PSC/EB and neuroectoderm cells, DE analysis was done to compare intermediate cells identified with CSS integration as described above with control cells. The comparisons to PSC/EB and neuroectoderm control cells were done separately. Here, control PSC/EB cells were defined as cells of PSC/EB samples with at least 40 of their 50 nearest neighbors being PSC/EB cells. Control neuroectoderm cells were defined in a similar way. Marker genes of the transition state were defined as genes with BH-corrected Wilcoxon rank sum test P<0.01, fold change > 1.5 and AUC > 0.6 in both comparison of transition-vs.-PSC/EB and transition-vs.-neuroectoderm.

### Calculation of CSS representation towards the scRNA-seq reference for query data

To calculate CSS representation of scRNA-seq data towards a reference, Spearman correlation coefficients were calculated between the transcriptomic profile of each query cell and average transcriptomic profiles of clusters in the reference samples. For each query cell, correlations to clusters of the same reference sample were normalized. The normalized similarities to clusters in different reference samples were then concatenated for the final representation.

To calculate CSS representation of scATAC-seq data towards a scRNA-seq reference, we firstly summarized peak accessibilities to gene activity scores for each cell. The detected peaks were annotated using the R package ChIPseeker [24], against the gene annotation of UCSC (hg19). For each cell, the proportion of detected genic peaks, defined as peaks annotated to be at the promoter, exonic or intronic region of genes, was calculated (denoted as pi for cell i). For each gene with at least ten genic peaks detected in the data set, its proportion of detected genic peaks at each cell was also calculated (denoted as pi,j for gene j in cell i). The gene activity score of gene j in cell i was then defined as the odds ratio pi,j / pi. CSS representation was then calculated for each cell against the reference sample clusters, by calculating, normalizing and concatenating the Spearman correlation coefficients between the gene activity profile of cells in the scATAC-seq data and the average transcriptomic profiles of the reference sample clusters.

## Supplementary Figures

**Supplementary Fig. 1.**
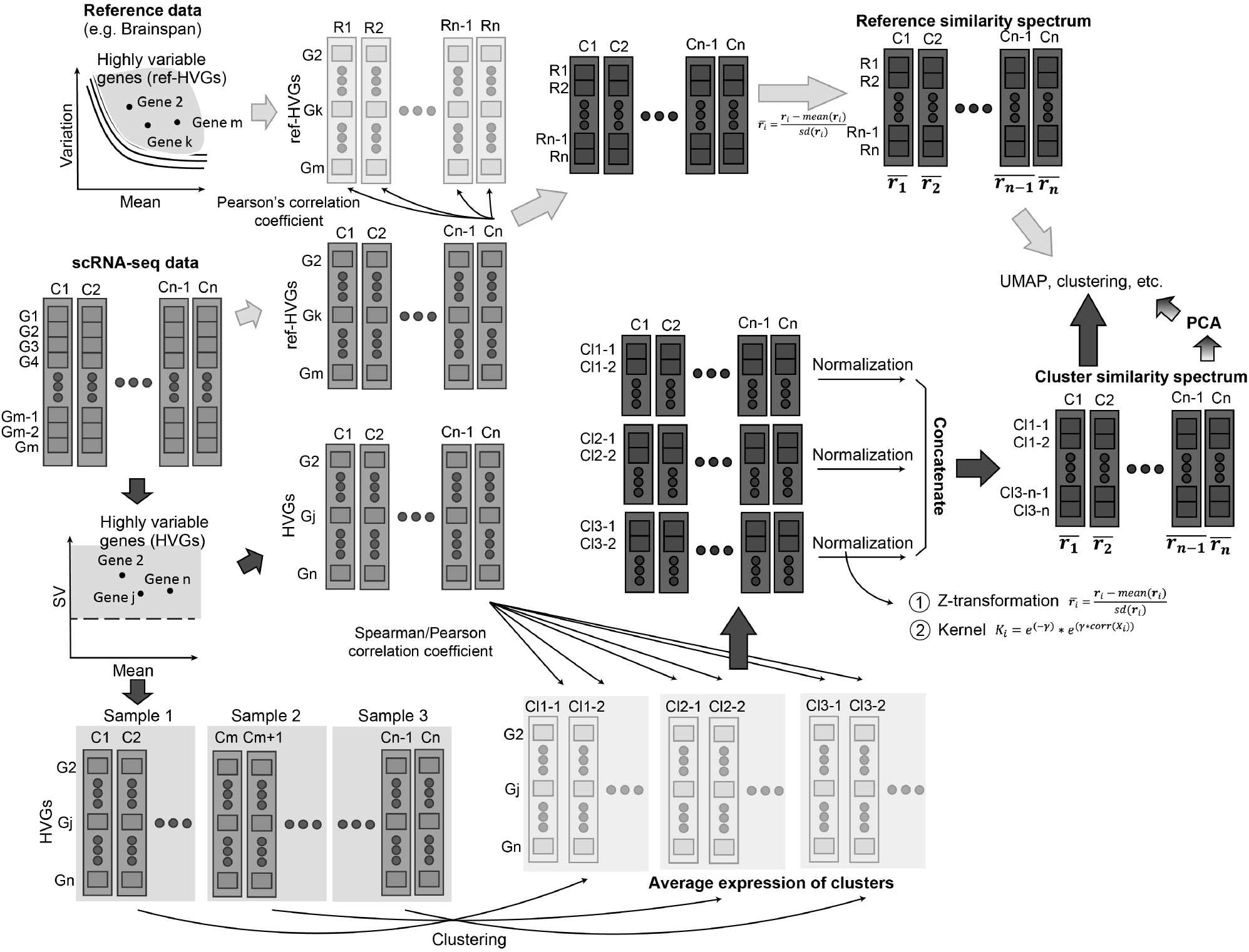
Detailed schematic of RSS and CSS representation calculation. To calculate RSS, highly variable genes in the reference data are first identified. Transcriptome similarity between each cell in the scRNA-seq data is calculated to each of reference samples across the identified genes. The resulting correlations of one cell are normalized using z-transform to obtain its RSS representation. To calculate CSS, clustering is applied to cells in each sample separately. Average transcriptome profiles of clusters in samples are calculated. Similarities between transcriptome of each cell and the cluster average transcriptome profiles are calculated. For each cell, similarities to clusters of one sample are normalized by z-transform or kernel probability transformation, with the normalized similarities to different samples concatenated to obtain its CSS representation. The CSS representation can be directly used as the input of the following analysis, or alternatively, an extra PCA is applied to the CSS representation to further condense the information and reduce the dimensionality.

**Supplementary Fig. 2.**
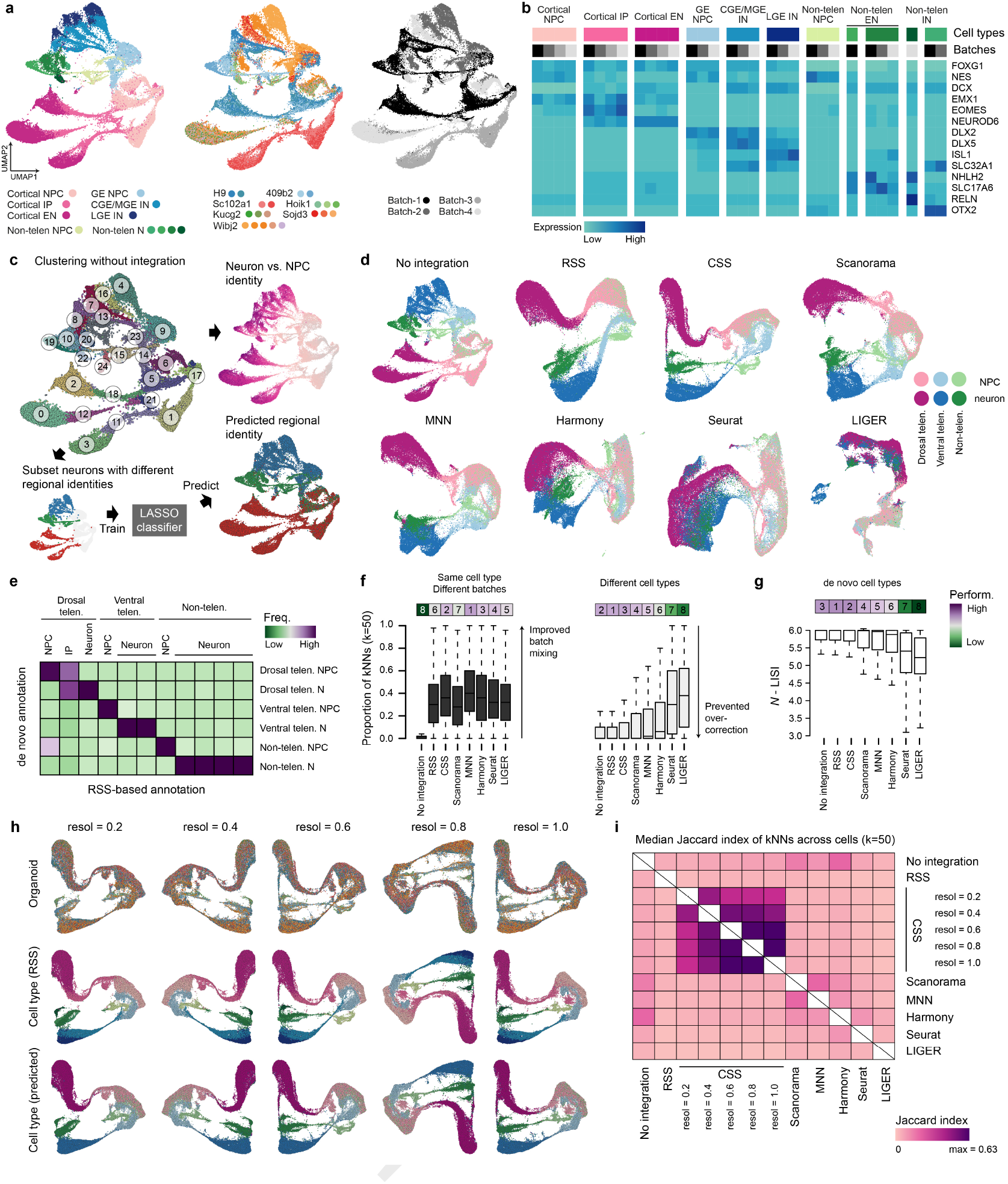
Cell type annotation of the cerebral organoid scRNA-seq data and CSS robustness to clustering of samples. (a) Joint analyzing scRNA-seq data of all the twenty cerebral organoids with no integration. The UMAP embedding is colored by RSS-based cell type annotation (left), organoids (middle), and experimental batches (right), respectively. (b) Average expression patterns of selected cell type markers in different cell types, with cells from organoids of different experimental batches separated. (c) Schematic of the model-based prediction of cell type. A random subset of cells in the dorsal/ventral/non-telencephalic neuron clusters were selected to train a prediction model of regional identities. The model was applied to all cells for their predicted regional identities. This information was combined with the NPC/neuron identity estimated by comparing expressions of NPC/neuron markers to get the predicted cell type annotation. (d) UMAP embeddings based on no integration or one of the seven integration methods, colored by the predicted cell type annotation. (e) Heatmap showing consistency between the RSS-derived annotation and the predicted cell type annotation. (f) Proportion of neighbors of each cell which (left) share the same predicted cell type but from a different batch, or (right) are of different predicted cell types. Neighbors are defined as 50 cells with the shortest Euclidean distances with the cell in PCA (no integration) or different integration spaces. Bars on top are colored by the median proportions, with the numbers showing ranks of different methods. (g) LISI-based scores of the model-based cell type annotation with no or different integration methods. Bars on top are colored by the median proportions, with the numbers showing ranks of different methods. (h) UMAP embeddings based on CSS with different clustering resolution used at the step of cell clustering for each sample separately. The embeddings are colored by organoids (upper row), RSS-derived annotation (middle row) and the predicted cell type annotation (bottom row). (i) Pairwise similarity between kNN graphs (k=50) generated using different integration methods with different parameters. The similarity between two kNN graphs is represented as the median Jaccard index of cell neighborhood in the two graphs across all cells.

**Supplementary Fig. 3.**
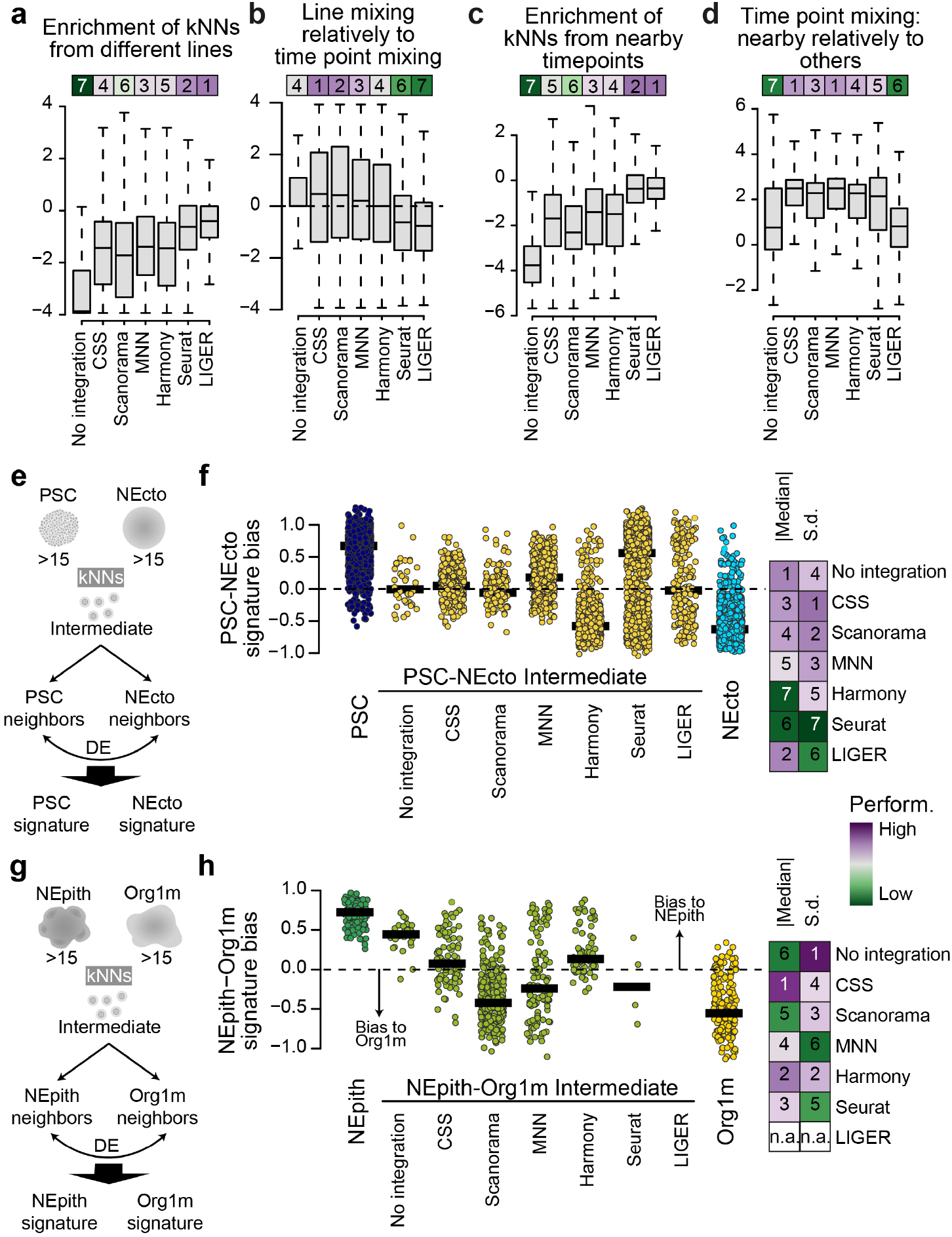
Benchmark of different integration methods on the time course cerebral organoid scRNA-seq data set. (a) For each integration method,boxplots show enrichment of neighboring cells from a different line in comparison to the global proportion of cells from different lines in the whole data set. An increased enrichment indicates improved mixing of cells from different lines. (b) Enrichment of neighboring cells from a different line relative to that of neighboring cells from a different time point. (c) For each integration method, boxplots show enrichment of neighboring cells from nearby time points, in comparison to the global proportion of cells from different time points in the whole data set. An increased enrichment suggests improved connectivity between cells from nearby time points. (d) Enrichment of neighboring cells from nearby time points relative to other time points. A higher relative enrichment suggests better resolving the temporal orders of the biological processes. Bars on top of (a-d) are colored by the median proportions, with the numbers showing ranks of different methods. (e) Schematic of identifying and benchmarking intermediate cells between PSC and neuroectoderm (NEcto) time points. (f) Benchmark of signature unbiasedness of identified PSC-NEcto intermediate cells. The scatter plot shows distributions of PSC-NEcto signature biases of PSC-NEcto intermediate cells identified based on different integration methods. Every dot represents one cell and the lines show the medians. The heatmap shows the summarized scores to rank different integration methods. A good set of intermediate cells is expected to concentrate across zero bias. (g-h) Similar to (e-f) but to identify and benchmark intermediate cells between neuroepithelium (NEpith) and one-month-old organoids (Org1m) time points.

**Supplementary Fig. 4.**
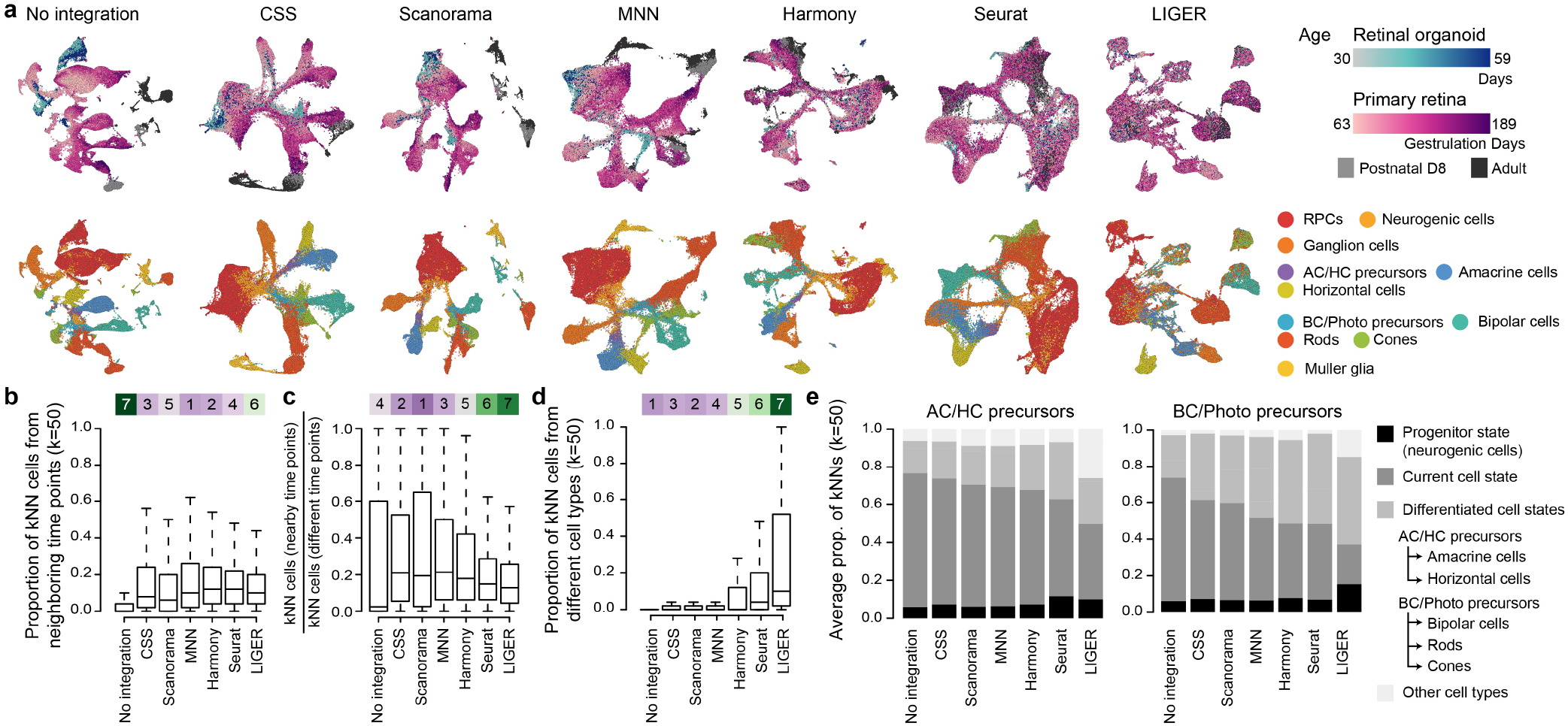
Benchmark of different integration methods on the time course human retinal organoid and primary retina scRNA-seq data set. (a) UMAP embeddings based on no integration or one of the six integration methods, colored by the sample ages (upper) and annotated cell types (lower). (b) Proportion of neighbors of each cell which were from samples at neighboring time points. (c) Proportion of neighbors which were from the neighboring time points among all neighbors which were from any different time point. (d) Proportion of neighbors annotated as different cell types. Neighbors are defined as 50 cells with the shortest Euclidean distances with the cell in PCA (no integration) or different integration spaces. Bars on top of (b-d) are colored by the median proportions, with the numbers showing ranks of different methods. (e) Stacked bar plots showing average proportions of cells annotated as the progenitor cell type (dark), same cell type (grey), differentiated cell type (lighgrey), and others (lightest grey), in the neighborhood of cells annotated as two groups of intermediate precursors: AC/HC precursors (left) and BC/Photo precursors (right), when no integration or different integration methods are used.

**Supplementary Fig. 5.**
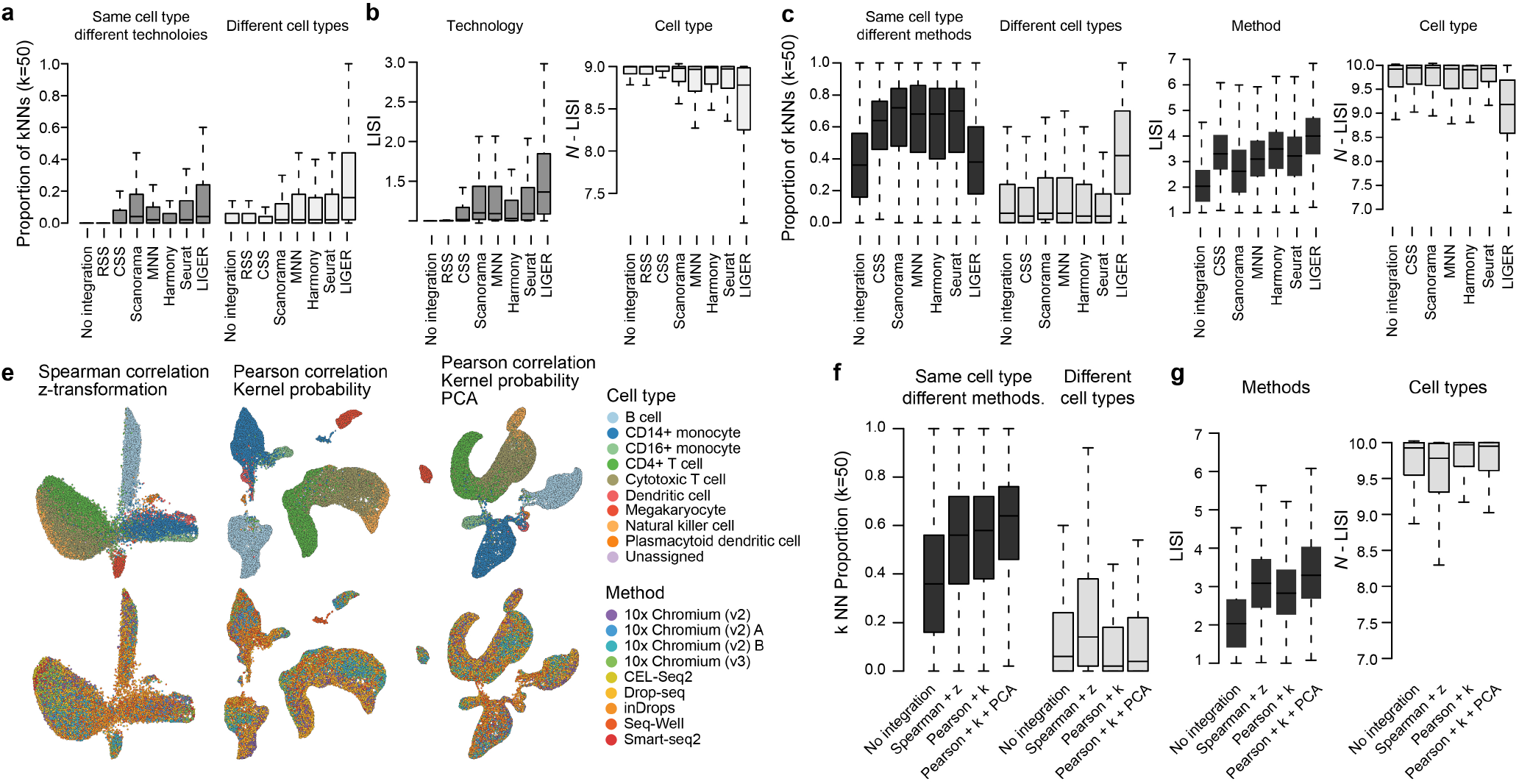
Benchmark of different integration methods on scRNA-seq technologies. (a-b) Benchmark of different integration methods on the cerebral organoid scRNA-seq data set with three different technologies. (a) Proportion of neighbors of each cell which (left) share the same cell type but measured by different technologies, or (right) are of different cell types. Neighbors are defined as 50 cells with the shortest Euclidean distances with the cell in PCA (no integration) or different integration spaces. (b) LISI-based scores of technologies (left) and cell type annotation (right) with no or different integration methods being used. A higher score indicates better performance. (c-d) Benchmark of different integration methods on the PBMC scRNA-seq data set measured by different methods. (c) Proportion of neighbors of each cell which (left) share the same cell type but measured by different methods, or (right) are of different cell types. Neighbors are defined as 50 cells with the shortest Euclidean distances with the cell in PCA (no integration) or different integration spaces. (d) LISI-based scores of methods (left) and cell type annotation (right) with no or different integration methods being used. A higher score indicates better performance. (e) CSS-based UMAP embeddings of the PBMC scRNA-seq data set with different CSS parameters. The embeddings are colored by annotated cell types (upper row) and methods (lower row). (f-g) Similar to (c), but to benchmark different CSS settings for the PBMC scRNA-seq data set.

